# Temperature alters *Culex tarsalis* West Nile virus vector competence, tissue bottlenecks, and transcriptional responses

**DOI:** 10.64898/2026.11.04.763972

**Authors:** Emily N Gallichotte, David C King, Briana Marsico, Emily A Fitzmeyer, Hunter A Ogg, Kate X Kimball, Nora Ebel, Corey L Campbell, Gregory D Ebel

## Abstract

West Nile virus is an arbovirus transmitted by *Culex* mosquitoes. Temperature is frequently the strongest abiotic factor impacting transmission in many virus-vector pairings, including WNV and *Culex*. However, it is unknown how temperature impacts WNV population bottlenecks and transcriptional responses to infection in *Culex tarsalis*. We found that elevated temperatures led to increased likelihood of transmission of rare virus variants due to increased vector competence and less stringent tissue barriers. Analysis of *Cx. tarsalis* transcriptional responses confirmed that WNV infection had minimal impacts on transcriptional activity. Temperature, however, induced significant changes to mosquito gene expression, whereas canonical immune gene expression minimally changed in response to infection. Thus, temperature impacts bottlenecks in the *Culex*-WNV system independent of known antiviral pathways, possibly by altering virus replication dynamics. While temperature had a dramatic impact on bottleneck stringency, this is likely independent of antiviral pathways and instead some other change to mosquito biology. Together, these results shed light on the relationships between temperature, virus evolutionary dynamics and antiviral immunity.

**IMPORTANCE:** West Nile virus is transmitted by *Culex* mosquitoes, with birds as their primary host, however it can cause severe disease in humans. The degree to which increasing temperatures may alter WNV infection in mosquitoes, or the mosquito’s ability to respond to both virus and temperature, is currently unclear. At increased temperatures, mosquitoes have higher transmission rates and levels of virus but also have decreased survival. Additionally, as the virus moves through a mosquito, it faces multiple bottlenecks which randomly reduce the genetic diversity of the virus population. We found that at hotter temperatures, these bottlenecks are weaker. Interestingly, the mosquito host response to infection at all temperatures was minimal, whereas they responded much stronger to changes to temperature, independent of infection. Together these results reveal that changes to temperatures alter mosquitoes-virus interactions, impacting the mosquito host response, survival, and ability to transmit virus.

## INTRODUCTION

For an arbovirus to be transmitted by a mosquito, it must first feed on a viremic animal. The virus must then systemically infect the mosquito, passing through a series of tissue ‘barriers’ before being expectorated along with mosquito saliva into the next host [1]. Each tissue barrier imposes a physiological bottleneck on virus populations, which dramatically and stochastically (i.e., independent of natural selection) restructure viral populations [2]. Like all RNA viruses, WNV exists in nature as a diverse swarm of genetically distinct virus genomes [2]. This genetic diversity is critical for virus fitness, maintaining efficient infection in both vertebrate and invertebrate hosts, and allowing adaptation to new and dynamic environments (temperatures, host species, etc.) [3]. This is achieved via mutations throughout the viral genome resulting in viruses with varying levels of fitness. Barcoded viruses are an important tool to study the genetic diversity in RNA virus quasispecies swarms without altering viral fitness [4]. We developed a fitness-neutral, clone-derived barcoded WNV (bcWNV) to mimic arbovirus quasispecies diversity [5, 6], and demonstrated mosquito tissue bottleneck stringency varies by species, with the most competent *Cx. tarsalis* having the weakest bottlenecks [6]. However, it is unknown how other variables, such as temperature, impact bottleneck stringency on WNV within *Cx. tarsalis*.

Several intrinsic (e.g., viral and mosquito genetics, microbiome, etc.) and extrinsic (e.g., temperature, humidity, etc.) factors impact vector competence for any vector-virus pairing. Climate uncertainty has spurred numerous studies focused on the impacts of temperature on mosquitoes. As ectotherms, mosquitoes are entirely dependent on the temperature of their environment, and temperature can have large impacts on habitat range [7, 8], life history traits (e.g., development, thermal tolerance, fecundity, etc.) [9, 10], and virus-mosquito interactions [11, 12]. Vector competence studies overwhelmingly show that across many vector-virus pairings, including WNV in multiple *Cx.* species [13–20], increased temperatures result in shorter extrinsic incubation period (EIP), increased infection, dissemination and/or transmission rates, and higher virus levels [11, 12]. It is unknown though, whether this phenotype is due to direct temperature impacts on the virus, modifications of the host response (e.g., anti-viral, heat-shock, etc.), a combination of both, or other undescribed mechanisms.

Unlike well-established invertebrate model organisms with fully annotated genomes (e.g., *Drosophila melanogaster*, *Caenorhabditis elegans*) there are relatively few studies evaluating comprehensive host transcriptional changes in mosquitoes. The majority have been performed in *Aedes* and *Anopheles*, primarily in response to blood-feeding [21–23], evaluating insecticide resistance [24–27], and infection [28–32]. Fewer studies have assessed how temperature impacts mosquito RNA expression and immunity. One study evaluated thermal variation on *An. stephensi* transcriptional responses [33], and two additional studies evaluated the impact of temperature on *Ae. aegypti* during chikungunya virus (CHIKV) and Zika (ZIKV) virus infection [34, 35]. Interestingly, both studies of infected *Ae. aegypti* saw minimal host transcriptional changes at increased temperatures compared to cooler temperatures [34, 35].

Compared to *Aedes* and *Anopheles*, relatively few studies have addressed transcriptional responses in *Culex*, likely due to fewer genetic resources [36]. The *Cx. tarsalis* genome was published in 2021, but significant portions need further annotation. While the *Cx. quinquefasciatus* genome was published in 2010, a newer chromosome-level assembly was only made available in 2024 [37–39]. Despite these limitations in genomic resources for *Culex* mosquitoes, some studies have been conducted to address their transcriptional responses to various infectious challenges and in different tissues, etc. [40–47]. Previously, we used single-cell RNA sequencing to construct a *Cx. tarsalis* midgut cell atlas and quantify cell-type specific host gene expression during WNV infection [46]. Unlike most *Aedes*-arboviruses pairings, which result in a robust host response, we found limited transcriptional responses following WNV infection, and minimal changes to expression of canonical immune pathway genes consistent with a prior study [40, 46]. How *Cx. tarsalis* transcriptional response is altered in response to changes in temperature, both with and without WNV infection, remains unknown.

We therefore sought to quantitatively evaluate the impact of altered temperature on *Cx. tarsalis* vector competence, tissue-associated bottlenecks, and their impact on viral population diversity and complexity, and host response to both temperature and infection. Using standard vector competence experiments and deep-sequencing (of both virus and host), we found that increased temperature weakens tissue bottlenecks, increasing both vector competence and altering diversity and complexity of viral populations. Additionally, while infection had a limited impact on *Cx. tarsalis* transcriptional response, temperature alone, and in combination with WNV infection, resulted in greater global changes to gene expression.

## RESULTS

### Impact of temperature on WNV-infected mosquito survival

*Cx. tarsalis* pupae were transferred from their standard rearing conditions of 26°C, to environmental temperatures of 22°C, 26°C or 30°C and ∼90% humidity (**SFig. 1**). Mosquitoes were fed either a mock or WNV-containing bloodmeal, survival measured daily, and on days 6, 12 and 18, a subset of mosquitoes was collected for mosquito and virus analyses (**SFig. 1**). At 22°C and 26°C, survival throughout the study duration was approximately 80% in mock and infected mosquitoes (**Fig. 1a**). At 30°C however, significantly more WNV-infected mosquitoes died (∼50%) compared to uninfected controls (∼27%) (**Fig. 1a, SFig. 2**). There was no difference in survival between mock mosquitoes at the three different temperatures. However, significantly more WNV-infected mosquitoes died at 30°C compared to infected mosquitoes at 22°C and 26°C (**Fig. 1b**, **SFig. 2**).

**Figure 1.**
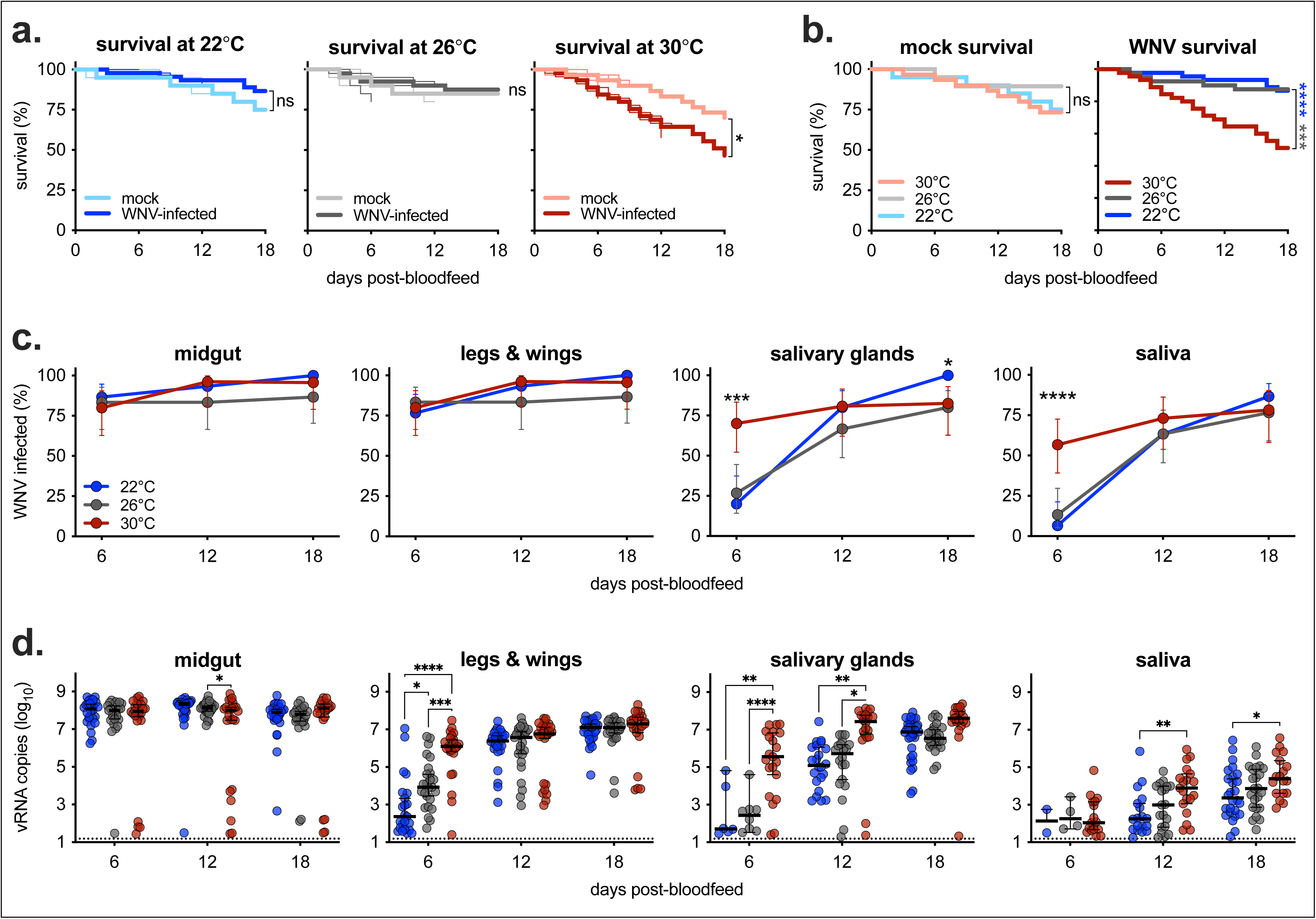
Higher temperature decreases *Cx. tarsalis* survival and increases WNV infection rates and virus levels. **a, b)** Survival was measured daily for mock (n=20-30/replicate), and WNV-infected (n=40-45/replicate) mosquitoes. Three biological replicates for each temperature and infection condition, ending on days 6, 12 and 18 (each replicate shown as thin line), with average survival plotted in heavy line. Log-rank (Mantel-Cox) test of averages between each condition/temperature, * p<0.05, *** p<0.001, **** p<0.0001. **c)** At days 6, 12 and 18 post-bloodfeed, midguts, legs and wings, salivary glands, and saliva were collected, RNA extracted, and the presence of WNV RNA detected via qRT-PCR (n=23-30 per group). Percent infected calculated as the number of positive samples out of the total number tested with Wilson/Brown 95% confidence intervals. Fisher’s exact test was performed for each time point, *p<0.05, ***p<0.001, ****p<0.0001. **d)** Samples with detectable WNV RNA are plotted (median ± 95% confidence intervals). Dotted line indicates limit of detection. Two-way ANOVA with Tukey’s multiple comparison test was performed between temperatures at each time point, *p<0.05, **p<0.01, ***p<0.001, ****p<0.0001.

### Increased vector competence of WNV at higher temperatures

Midgut, legs and wings, salivary glands, and saliva from each temperature at each time point were assayed for the presence of viral RNA (vRNA) via qRT-PCR (**Fig. 1c**, **d**). Midgut, and leg and wing infection was comparable across all temperatures at all days post-bloodfeed (**Fig. 1c**). However, mosquitoes at 30°C had significantly more WNV-positive salivary glands and saliva samples at day 6 compared to 22°C and 26°C, with similar proportions infected by days 12 and 18 (**Fig. 1c**). Consonant with higher proportions of infected mosquitoes at 30°C, mosquitoes at this temperature had significantly higher levels of vRNA in legs and wings, salivary glands and saliva at multiple time points compared to 22°C and 26°C (**Fig. 1d**).

### Higher temperature results in weaker tissue bottlenecks

The WNV barcode region of a subset of mosquitoes from each temperature and time point were selected for analysis of the WNV barcode region (**SFig. 3**). Total barcode count results (which quantify the overall amount of virus genome in a given sample) were comparable to qRT-PCR-based quantification of viral load (**Fig. 1d**), with significantly more total viral barcodes in legs and wings, salivary glands, and saliva at 30°C (**Fig 2a**). In the midgut, the number of unique barcodes (which measures virus population richness) and population complexity (evenness of barcodes) decreased as temperatures increased, with highest richness and complexity at 22°C (**Fig. 2b**, **c**). In other sample types, diversity and complexity varied by temperature and day post-bloodfeed with few consistent trends by temperature (**Fig. 2b, c**)

**Figure 2.**
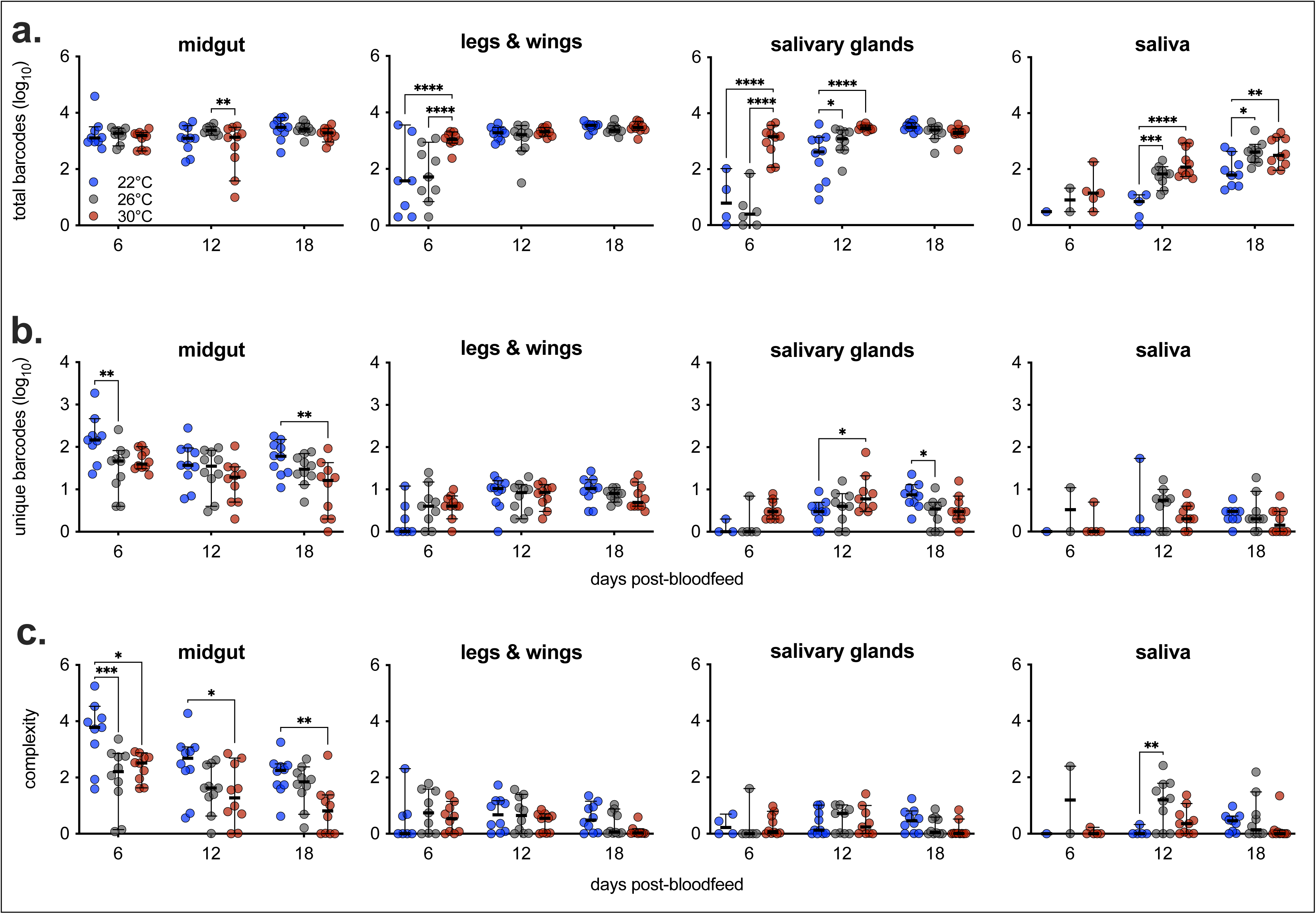
WNV barcode counts, richness and complexity varies by temperature and mosquito sample. **a)** Total barcodes, **b)** unique barcodes and **c)** complexity from each of the mosquito samples at the three experimental temperatures (22°C, 26°C and 30°C) (median ± 95% confidence intervals). With each sample type, two-way ANOVA with Tukey’s multiple comparisons test across temperatures within time-points, *p<0.05, **p<0.01, ***p<0.005, ****p<0.0001.

We next quantified the stringency of tissue bottlenecks by calculating the relative diversity through each tissue barrier (i.e., percent of unique barcodes present in the legs and wings that are also detected in the salivary glands at a given timepoint). Despite fewer unique barcodes in midguts with increasing temperatures (**Fig. 2b**), at 30°C more unique midgut-detected barcodes were also detected in legs and wings (**Fig. 3a**). The salivary gland infection barrier alone (legs and wings to salivary glands), and combined midgut and salivary gland infection bottlenecks (midgut to salivary glands) were also weaker at hotter temperatures, with higher average rates of relative diversity (**Fig. 3a**). With all bottlenecks combined (midgut to saliva), there was a trend of increased relative diversity with increasing temperatures (**Fig. 3a**).

**Figure 3.**
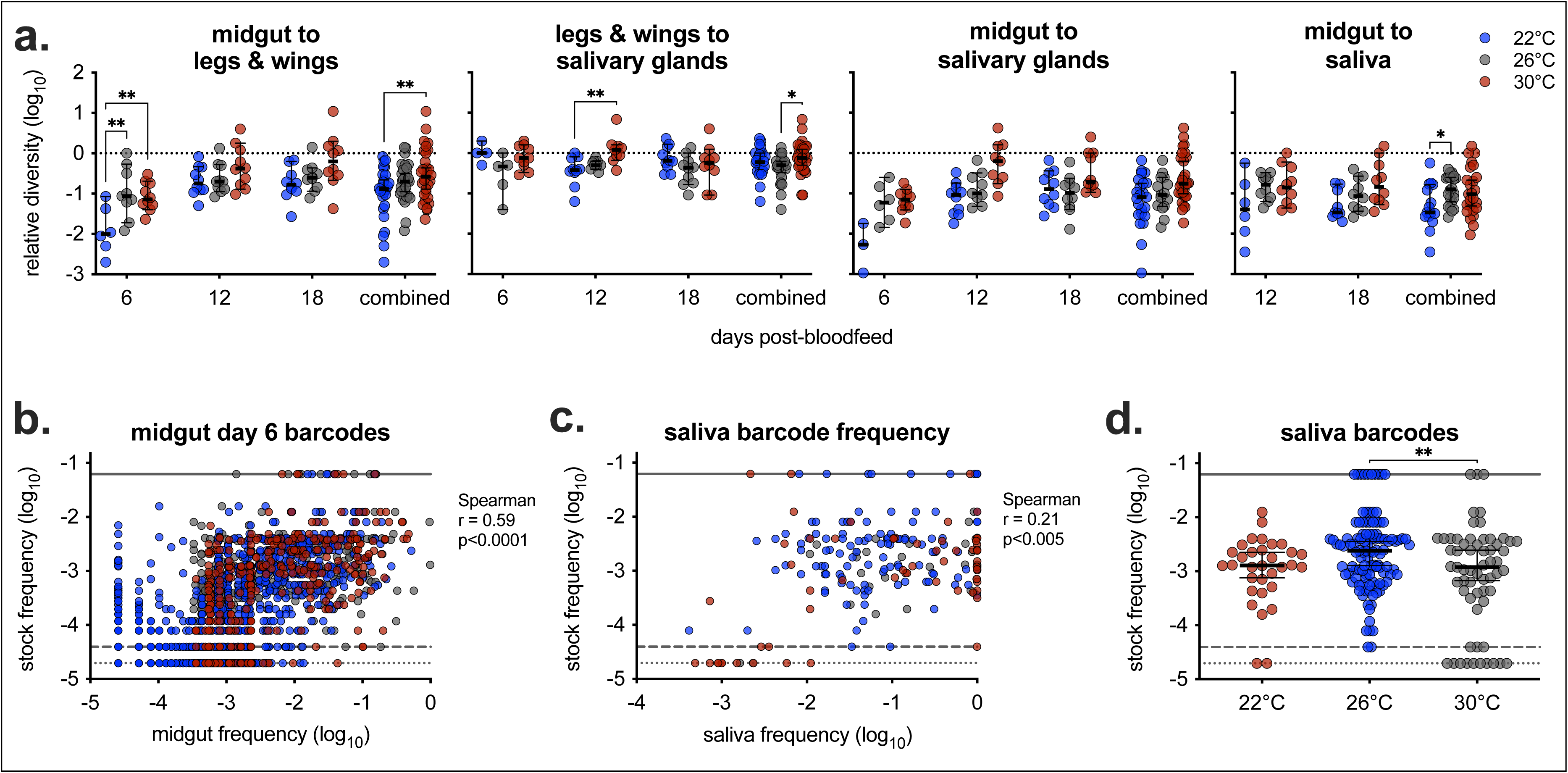
Mosquito tissue bottlenecks are weaker, and rarer barcodes are transmitted at hotter temperatures. **a)** Relative diversity of barcode populations across bottlenecks by temperature (22°C, 26°C and 30°C) and time point (individually, and time points combined) (median ± 95% confidence intervals). Relative diversity calculated as diversity (number of unique barcodes) post-bottleneck (e.g., legs and wings) divided by pre-bottleneck (e.g., midgut). Relative diversity of 0 log_10_ (dashed line) equals same diversity of virus populations pre-and post- bottleneck. For each bottleneck, two-way ANOVA with Tukey’s multiple comparisons test across temperatures within time-points, *p<0.05, **p<0.01. **b)** The frequency of all barcodes detected in the midgut at day 6 post-bloodfeed or **c)** saliva barcodes (all time points) for each temperature was compared to the corresponding frequency of that barcode in the stock virus (as described in Supplemental Figure 4). **d)** The stock frequency of all barcodes detected in the saliva (regardless of frequency) was compared by experimental temperature by one-way ANOVA with Tukey’s multiple comparisons test, **p<0.01 (median ± 95% confidence intervals). **b-d**) Solid line represents the most dominant barcode in the stock (6% of total reads), dashed line represents the frequency of barcodes only detected one time (0.004%) and dotted line represents barcodes not detected in stock (1/2 the frequency of those detected one time, 0.002%).

We next compared the frequency of barcodes detected in the midgut and saliva, to the frequency of that barcode in the virus stock (**SFig. 4a-d**). The frequency of barcodes in each of the bloodmeals was highly correlated to the barcode frequency in the stock virus (**SFig. 4e**). Not surprisingly, the frequency of barcodes in the midgut at day 6 was highly correlated to the stock frequency (**Fig. 3b**). However, frequency of barcodes detected in the saliva were poorly correlated (r = 0.21) to the stock barcode frequency (**Fig. 3c**). On average, mosquitoes at 26°C had virus in their saliva that was at significantly higher stock frequencies than those at 30°C, which also transmitted the greatest number of rare barcodes (those never detected in the stock) (**Fig. 3d**).

### Mosquito host response changes in response to temperatures

RNA-sequencing (RNAseq) analyses were conducted on mock and WNV-infected pools of midguts six days post-bloodfeed held at 22°C, 26°C, and 30°C (**SFig. 5**). We first compared infected to mock mosquitoes held at each temperature. We identified fifty-eight of 11,001 (0.527%) differentially expressed genes (DEGs) between WNV-infected and mock mosquitoes held at 22°C, whereas there were no DEGs comparing infection groups held at 26°C, and only one (0.009%) at 30°C. (**Fig. 4a**, **Table 1**). We then assessed differential gene expression among uninfected mosquitoes held at different temperatures to determine mosquito transcriptional responses to different environmental temperatures. There were nineteen (0.17%) DEGs between mock mosquitoes held at 22°C compared with 26°C; 273 (2.482%) between 26 and 30°C; and 220 (2.000%) between at 22°C and 30°C **(Fig. 4b, Table 1)**. Next, we assessed transcriptional responses to temperature in the presence of WNV infection, which overall resulted in substantially fewer changes. There were 54 (0.491%) DEGs in WNV-infected mosquitoes held at 22°C compared with 26°C; 26 (0.236%) between 26°C and 30°C; and 11 (0.100%) between 22°C and 30°C. (**Fig. 4c, Table 1**).

**Figure 4.**
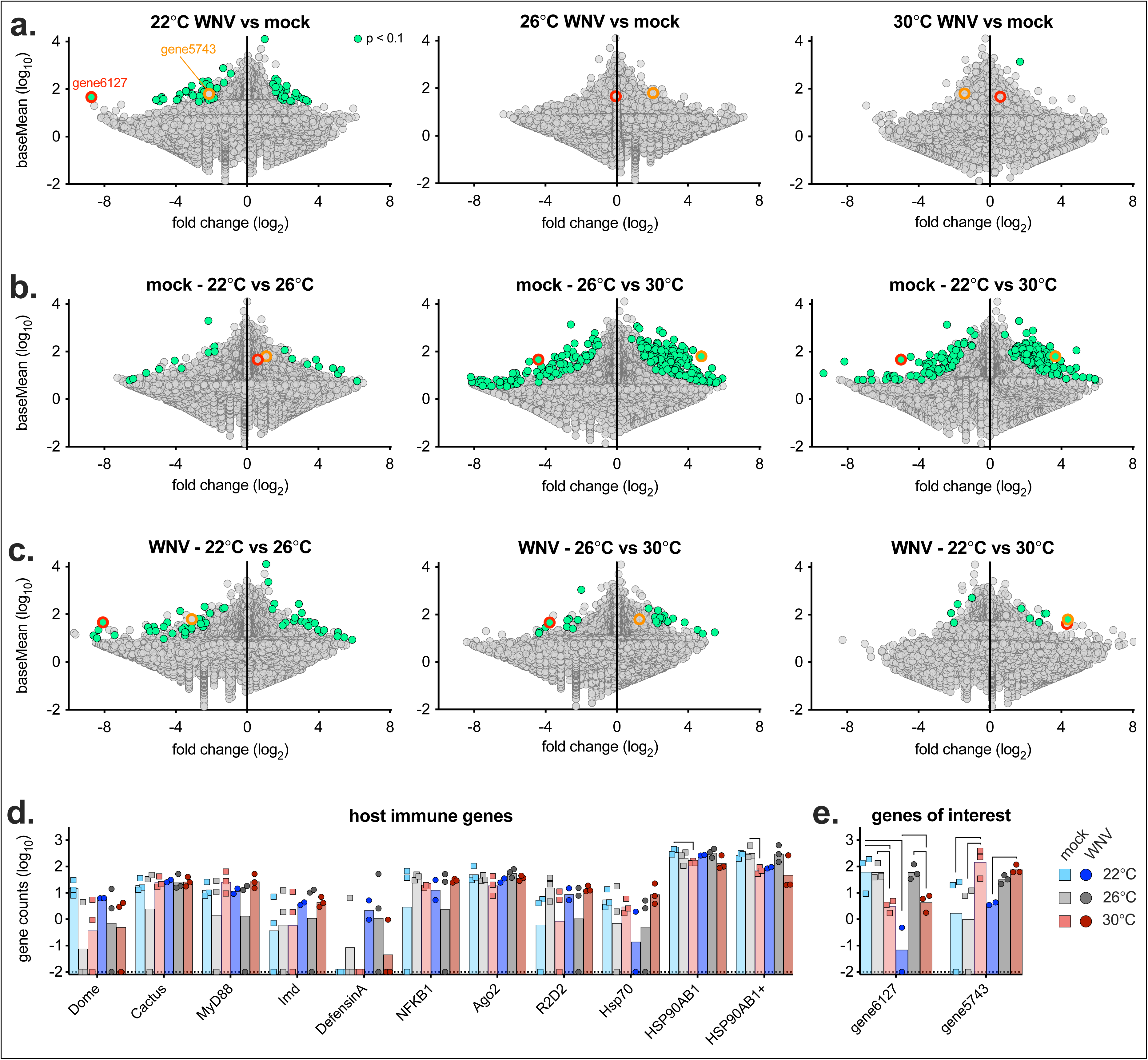
Mosquito RNA response changes more significantly by altered temperature than by WNV infection. MA plots comparing fold change and expression (baseMean) of 11,001 *Cx. tarsalis* genes by **a)** infection within a temperature, or across experimental temperatures (22°C, 26°C and 30°C) with **b)** mock and **c)** WNV-infected mosquitoes. Genes with adjusted p-value < 0.1 are colored green. Two genes of interest with significantly different fold-changes across multiple conditions are colored with a red (gene6127) and orange (gene5743) outline. Gene counts of a panel of **d)** host immune genes and **e)** genes of interest (gene6127 and gene5743). Samples with no counts detected plotted at -2 (0.01 counts, limit of detection, dashed line), bars show mean of replicates. Brackets show significant (adjusted p-value < 0.1) fold-change from RNAseq analyses (Fig. 4a**-c**).

**Table 1.** Differentially expressed genes.

| **comparison** | **DEGs**<br>**total (# up, # down)** | **% DEGs** |
|---|---|---|
| 22°C, mock vs WNV | 58 (25, 33) | 0.527% |
| 26°C, mock vs WNV | 0 (0, 0) | 0.000% |
| 30°C, mock vs WNV | 1 (1, 0) | 0.009% |
| mock, 22°C vs 26°C | 19 (9, 10) | 0.173% |
| mock, 26°C vs 30°C | 273 (185, 88) | 2.482% |
| mock, 22°C vs 30°C | 220 (137, 83) | 2.000% |
| WNV, 22°C vs 26°C | 54 (23, 31) | 0.491% |
| WNV, 26°C vs 30°C | 26 (16, 10) | 0.236% |
| WNV, 22°C vs 30°C | 11 (7, 4) | 0.100% |

DEG = differentially expressed gene
% DEG based out of 11,001 detected total genes

Due to the low overall number of DEGs, gene set enrichment analysis failed to identify any pathways that were consistently significantly up- or downregulated in any of the virus-temperature classifications (data not shown). Therefore, we used the proportions of DEGs, regardless of function, as a proxy of transcriptional activation in response to infection and temperature (**SFig. 6**). There were significantly more DEGs in response to WNV infection at 22°C than 26°C or 30°C (**SFig. 6a**). Uninfected mosquitoes held at 30°C had significantly more DEGs than those held at either of the lower temperatures (**SFig. 6b**). Interestingly, this result was the opposite in WNV-infected mosquitoes, where there were significantly more DEGs between 22°C and 26°C, compared to mosquitoes held at 30°C (**SFig. 6c**). We next assessed the role of WNV infection on the proportion of DEGs for each temperature comparison. While relatively few genes change regardless of WNV infection between mosquitoes held at 22°C compared to 26°C, when temperatures are increased to 30°C, WNV-infection significantly reduces the number of DEGs compared to mock mosquitoes (**SFig. 6c**).

We finally compared gene expression levels of a panel of host immune and heat-shock genes. Expression of canonical immune genes (e.g., JAK-STAT and RNAi pathways, heat-shock, etc.) was generally not consistently upregulated across samples and conditions (**Fig. 4d**). Two genes changed significantly in response to both infection and temperature in multiple comparisons (**Fig. 4e**). While variable, gene6127 expression levels generally decreased in response to increased temperature, whereas gene5743 expression increased with increased temperatures (**Fig. 4d**).

### Gene5743 increases in response to temperature and infection in the midgut and salivary glands

We validated the expression changes of gene6127 and gene5743 of the same midgut pools used for RNAseq using qRT-PCR (**Fig. 5a**). Gene6127 levels were more comparable across conditions; however gene5743 expression was significantly increased at higher temperatures in both mock and WNV-infected mosquitoes (**Fig. 5a**). We then compared gene6127 expression in the midguts at day six of the corresponding full set of mock and WNV-infected mosquitoes from the larger study (**Fig. 1**). There was a bimodal expression of gene6127 across both conditions and all temperatures, with no significant differences (**Fig. 5b**). However, when we divided the results into high and low expression, WNV-infected mosquitoes had higher expression than mock at the corresponding temperatures, and within WNV infection, significantly lower expression at 30°C (**Fig. 5b**). We then quantified gene5743 expression in midguts in all conditions at all time points and saw that relative to mock 26°C at each time point, gene5743 expression increases with temperature in both mock and WNV-infected mosquitoes (**Fig. 5c**). Additionally, when comparing all conditions relative to day 0 at 26°C, gene5743 increases in response to bloodfeed, in a temperature-dependent manner (**Fig. 5d**). We then quantified gene5743 expression in salivary glands, and saw similar trends, with expression generally increasing in response to temperature and WNV-infection, especially at later time points (**Fig. 5e-f**).

**Figure 5.**
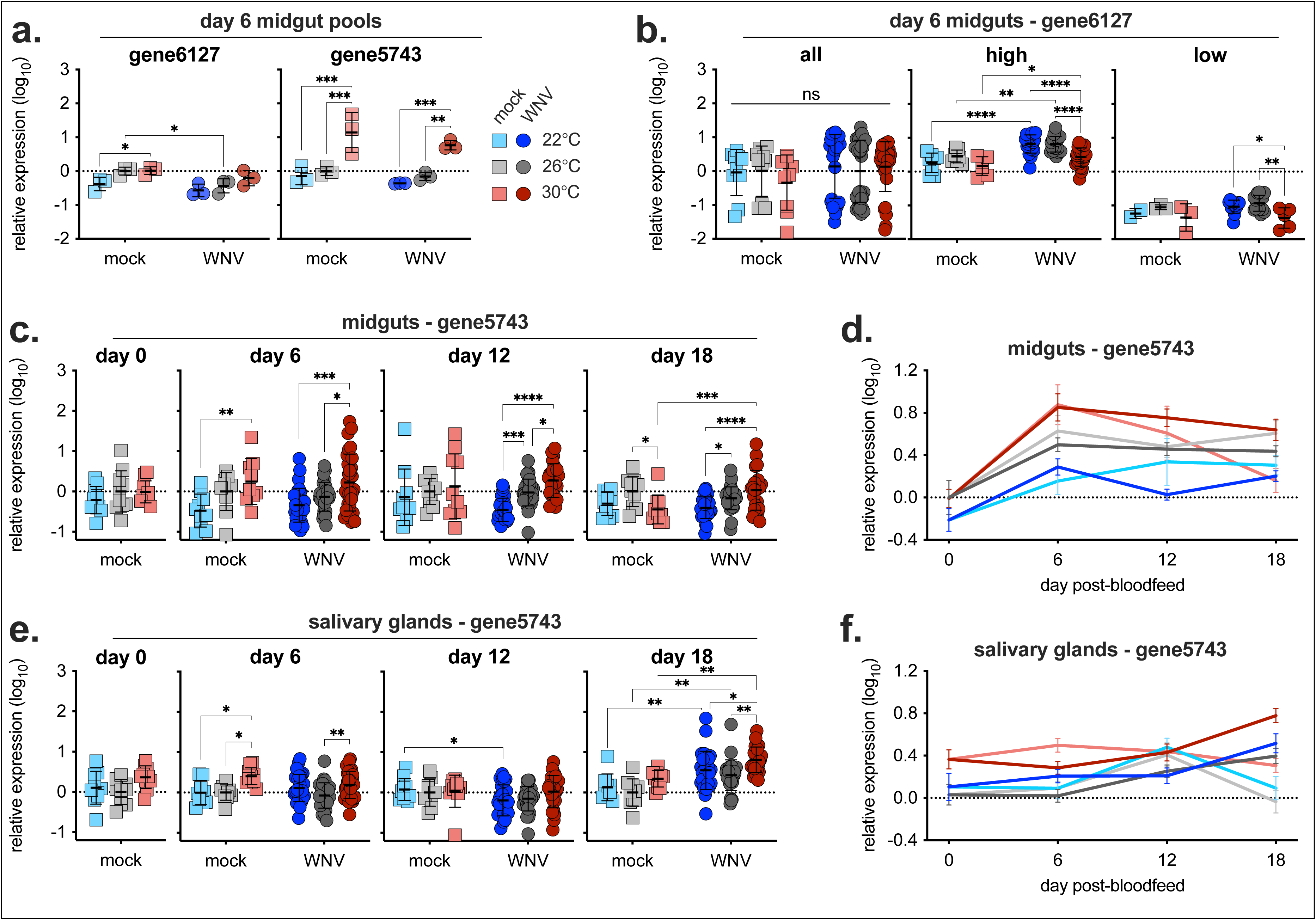
*Cx. tarsalis* gene5743 expression increases in the midgut and salivary glands at increased temperatures. Gene6127 and gene5743 expression levels were quantified via qRT-PCR. **a)** Relative expression in the same samples used in RNAseq analyses (three pools of six midguts each of day 6 mock and WNV-infected mosquitoes at 22°C, 26°C and 30°C). **b, c, e)** Relative expression in experimental mock and WNV-infected midgut and salivary gland samples (Fig. 1). **a, b, c, e)** Within each time point, relative expression normalized to housekeeping gene RPL37 and mock 26°C (mean ± standard deviation). Two-way ANOVA with Tukey’s multiple comparisons test across temperatures within condition and across condition within temperatures, *p<0.05, **p<0.01, ***p<0.001, ****p<0.0001. **d, f)** Relative expression normalized to housekeeping gene RPL37 and day 0 mock 26°C (mean ± SEM). Dashed line represents a relative expression of 0 (no change).

## DISCUSSION

While many studies have evaluated the role of temperature on arbovirus-mosquito vector competence outcomes, few have determined the impact to viral population structure and overall genetic diversity. Additionally, while the majority of studies show increased vector competence across many virus-vector pairings, it’s unknown whether this is due to temperature impacts on the virus directly (e.g., more efficient viral replication, packaging, etc.) or due to changes to the host which may improve infection (e.g., dampened anti-viral responses). In this study, we assessed *Cx. tarsalis* vector competence, tissue bottlenecks and impacts on viral populations, and host transcriptional changes in response to both WNV infection and temperature.

For many orthoflavivirus-mosquito pairings, infection has no fitness cost on survival, however, other life history traits may be impacted depending on the virus-vector pairing [48]. For WNV, it has been shown that infection has minimal negative impact on survival in *Cx. tarsalis*, *Cx. pipiens* or *Cx. quinquefasciatus* at standard temperatures, however some other outcomes including egg number, egg hatch rate and blood-feeding are altered by infection status [49–51]. Consistent with these reports, we found WNV infection had no significant impact on *Cx. tarsalis* survival at standard and cooler temperatures. Interestingly, WNV-infection significantly reduced survival at high temperatures (30°C), similar to as was seen in a WNV-infected colony of *Cx. quinquefasciatus* at 31°C [51]. This temperature is well within thermal limits for *Cx. tarsalis* [52]. This finding was consistent across two independent experiments. The mechanism causing increased mortality with infection at higher temperatures is unknown, but may be driven by virus-induced pathology or increased apoptosis of infected cells, as has been seen in WNV and other viruses [53]. Consistent with many other WNV-*Culex* studies [14–20], increased temperature resulted in increased dissemination and transmission rates, significantly higher levels of virus, and a shortened extrinsic incubation period (EIP). While mosquitoes held at increased temperature were capable of transmitting virus earlier, their decreased survival may limit their ability to transmit to a new host, potentially resulting in a similar overall vectorial capacity compared to standard temperatures [54]. Indeed, mathematical modeling found WNV transmission is optimized between 24–25°C, not hotter temperatures, despite the shorter EIP [55].

It is well established that as an arbovirus moves through its vector, the virus population is stochastically reduced via tissue (midgut and salivary gland) bottlenecks [2, 6, 56, 57]. The stringency of these bottlenecks can vary by mosquito species and strain [6, 56]. We found that temperature can strongly impact these as well. For example, at cooler temperatures, more complex populations with more unique barcodes were detected in the midgut. At cooler temperatures, mosquitoes take both larger bloodmeals [58, 59], and digest them slower, potentially allowing more time for viruses present in the bloodmeal to establish infection in midgut cells. Conversely, for subsequent bottlenecks (midgut escape, and salivary gland infection and escape), increased temperatures resulted in weaker bottlenecks, with a greater fraction of unique barcodes bypassing each barrier. Despite differences in overall vector competence and bottleneck stringency, rare barcodes were transmitted in the saliva at all experimental temperatures, however mosquitoes held at 30°C on average transmitted rarer barcodes. Additionally, temperature is known to impact virus mutation rates, and increased temperatures has been shown to increase Zika virus diversification in *Aedes* mosquitoes [60–62]. This increased diversification coupled with weaker bottlenecks and transmission of more rare variants at hotter temperature, makes it likely that warmer temperatures may facilitate the generation and emergence of rare variants.

Mosquitoes have several well-characterized immune pathways that are thought to play critical roles in responding to and controlling viral infections [63]. Interestingly, we saw limited overall host transcriptional responses, including in immune pathways, to WNV in *Cx. tarsalis*. Unlike many *Aedes*-virus pairings, which result in robust up-regulation of many canonical immunity genes [32, 64, 65], it appears WNV does not elicit a similar response in midguts of *Cx. tarsalis* or *Cx. quinquefasciatus* [40, 45, 46]. Importantly, others found strong induction of host immune pathways (e.g., Toll, Imd, etc.) in *Cx. tarsalis*, Cx*. quinquefasciatus* and *Cx. pipiens* following avian malaria or Rift Valley fever virus infection, demonstrating these pathways are responsive to infection with other pathogens [44, 47, 66]. It has been hypothesized that the non-responsiveness of these canonical immunity genes in response to WNV in *Culex* may reflect the persistent nature of arboviruses in their susceptible vector [40], however this does not explain the strong transcriptional response to many arboviruses in non-*Culex* species. Differences in immune gene activation and lack thereof depending on virus-vector pairing – and other likely important experimental conditions (e.g., time point, mosquito tissue, etc.) – may reflect the delicate interplay and co-evolution between viruses and their vectors, with mosquitoes mounting antiviral responses to control infection, and viruses developing strategies to suppress these host responses [67]. Additional research is needed to better define these relationships, especially using field-caught or more recently established mosquito populations, and contemporary viruses that reflect the infections occurring nature.

While we observed limited host transcriptional response to WNV infection, we saw a more dramatic changes in response to altered temperature, especially in uninfected mosquitoes similarly to as has been shown with other mosquito species [62]. Many *Culex* traits respond strongly and consistently to temperature, with some, like blood-feeding and fecundity, peaking around 24°C [55, 68]. Additionally, the thermal optimum for WNV transmission by *Cx. tarsalis* is modeled to be 23.9°C with a thermal breadth of 16.3-26.7°C [55]. Therefore, it is not surprising that in uninfected mosquitoes, we observed fewer transcriptional changes between 24°C and 28°C, as these are both within and near the thermal breadth. Conversely, mosquitoes held at 30°C are far outside their thermal breadth and nearing their thermal maximum (32°C) [55], likely physiologically stressing the mosquito, leading to the larger number of significantly differentially expressed genes. Many viruses, including arboviruses, induce host shut off as a strategy to enhance their replication and prevent anti-viral responses [69, 70]. Consistent with this, WNV infection appears to dampen the overall transcriptional response to temperature, resulting in fewer genes with differential expression. Together these results reveal that both infection and changes to temperature can dramatically alter host response both independently and together. Future analyses should investigate the function and roles of these differentially expressed genes and in response to both infection and temperature.

Little is known regarding the function and role of gene5743. Its homolog in other *Culex* and *Aedes* spp. encode predicted a MAP/microtubule affinity-regulating kinase and serine/threonine-protein kinases. Despite first identifying differential expression of gene5743 based on midguts at day 6 post-infection, its expression remained increased in response to temperature through 18 days post infection. In the salivary glands, while there were subtle changes in gene expression as early as 6 days post-infection, there were more dramatic increases in gene expression in response to both temperature and WNV infection at day 18 post-infection, when the majority of salivary glands were infected. The role this gene plays in both tissue types, and potentially others such as brain and ovaries, remains unknown.

Together, these results demonstrate that increased temperature dramatically impacts *Cx. tarsalis* mosquitoes, which in turn alters WNV vector competence and viral population structure and diversity within the mosquito. How these results compare with other important *Culex* vectors of WNV (e.g., *Cx. pipiens*) and related viruses of medical importance (e.g., St. Louis encephalitis virus) warrants future investigation.

## METHODS

### Viruses and mosquitoes

Barcoded West Nile virus (bcWNV) was generated and used as previously described [5, 6]. bcWNV contains random, degenerate synonymous nucleotides at the third nucleotide position of 11 consecutive codons within the NS4b coding sequence, resulting in 4^11^ possible unique barcodes [5, 6]. bcWNV for these experiments was used after two passages on Vero cells. The *Cx. tarsalis* colony was established from a colony by WK Reisen collected in California [71]. All mosquitoes were maintained at 26°C with a 16:8 photoperiod and 70–80% relative humidity. Water and sugar were provided *ad libitum*, and larvae were raised on powdered fish food.

### Mosquito infections and sample collection

Pupae were transferred into the experimental temperature (22°C, 26°C or 30°C) with ∼90% relative humidity. Five to seven days post-eclosion, sugar and water were removed, and ∼24 hours later, mosquitoes were fed a warmed (37°C) bloodmeal containing 2x10^6^ PFU/ml bcWNV or DMEM in defibrinated calf’s blood (final bcWNV concentration of 1x10^6^), using water-jacketed glass feeders sealed with hog’s gut. After one hour of feeding, mosquitoes were cold-anesthetized, and engorged females were sorted and returned to their experimental temperature. Bloodfed mosquitoes were held for 6, 12, or 18 days, with mortality measured daily. On the indicated day, mosquitoes were cold-anesthetized, legs and wings removed and salivary secretions collected using capillary tubes containing immersion oil for 30 minutes. Midguts and salivary glands were then dissected. Midguts, legs and wings, and salivary glands were placed into tubes containing a steel ball bearing and mosquito diluent (PBS, 20% FBS, 100 IU/mL penicillin, 100μg/ml streptomycin, 50µg/mL gentamicin, and 2.5µg/mL amphotericin B), homogenized, then centrifuged. Saliva-containing capillary tubes were placed into tubes containing mosquito diluent and centrifuged to expel saliva into the media. Samples were stored at -80°C.

### RNA extractions and qRT-PCR

Samples were thawed and RNA extracted using the Omega Bio-tek Mag-Bind® Viral DNA/RNA 96 Kit on a KingFisher™ Flex Purification System following manufacturers protocol. qRT-PCR was performed using the EXPRESS One-Step qRT-PCR kit following manufacturer’s instructions and primers and probes (**STable 1**). Primers and probes for mosquito genes were designed to span exon junctions. WNV RNA standards were used to generate copy numbers as previously described [72]. Delta delta CT method was used to analyzed host gene expression by normalizing to RPL37 levels, then mock mosquitoes (the time point used for normalization is specified in figure legends).

### Barcode WNV library preparation

Ten mosquitoes from each temperature and time point were selected for next generation sequencing of a 204bp amplicon containing the barcode region (**SFig. 3A**). Libraries were prepared as previously described [5, 6]. All primers used in the library preparation are provided (**STable 2 and 3**). Briefly, cDNA was generated using a WNV-specific primer and SuperScript IV reverse transcriptase. Reaction products were purified using 1X AmpureXP beads on a KingFisher Flex Purification System. Amplification of the barcode region was performed using KAPA HiFi HotStart ReadyMix and first round PCR primers, then purified as described above at a 0.7X concentration. Second round PCR was performed using KAPA HiFi HotStart ReadyMix and second round PCR primers which contained Illumina i5 and i7 10bp indexes, then purified up at a 0.7X concentration. DNA concentration of samples were quantified using Quant-iT dsDNA High Sensitivity Assay Kit, pooled at 2ng/sample then purified and concentrated at a 1.4X concentration.

### Mosquito RNA library preparation

Mosquitoes were reared and bloodfed as described above. On day 6 post-bloodfeed, six midguts per condition (mock and WNV, 22°C, 26°C or 30°C) were pooled and homogenized, then extracted in Tri-reagent using the Zymo Research Direct-zol RNA Miniprep Plus according to manufacturer’s instructions. Mosquito ribosomal RNA was depleted as described in Fauver et al. [73]. Briefly, RNA was DNase treated using TURBO DNase, then purified using RNAClean XP beads. Samples were incubated with depletion probes, and cDNA generated using AMV reverse transcriptase. Samples were treated with RNase H and DNase and purified using RNAClean XP beads. Ribosomal RNA depletion was confirmed on a TapeStation using a High Sensitivity RNA ScreenTape, and qRT-PCR for 18S gene. RNA sequencing libraries were then generated using the NEBNext Ultra II RNA Library Prep Kit for Illumina and NEBNext Multiple Oligos for Illumina (Dual Index Primers Set 1) following manufacturer’s instructions (**STable 4**).

### Sequencing and analysis

Both libraries were loaded at a concentration of 10nM on an Illumina NovaSeq 6000 with paired-end 2x150bp reads. bcWNV sequences were analyzed as previously described [6]. Briefly, data were demultiplexed, trimmed, paired-end reads merged, aligned to WNV genome, grouped by unique molecular identified and consensus sequences were generated [6]. Barcode abundance was determined, and population richness and complexity (Shannon’s index) were calculated as previously described (https://github.com/fitz-meyer/bcWNV_analysis). Relative diversity was calculated as diversity (number of unique barcodes) post-bottleneck (e.g., legs and wings) divided by pre-bottleneck (e.g., midgut).

RNAseq analyses were performed similarly as was previously described [74]. Sequencing reads were trimmed and filtered using Fastp version 0.23.4 [75]. Duplicate reads were retained. A reference transcriptome was created using the CtarK1 genome with exon annotations and canonical splice sites calculated using Hisat2 2.2.1 [76]. Reads were then aligned to this *de novo* transcriptome using splice-aware alignment. Only paired reads which aligned concordantly were retained. Gene-level aggregation was performed using HTseq-Count version 2.0.5 with the aforementioned annotations. Reads were compared against both strands.

Differential expression analysis was performed in DESeq2 [77] version 1.48.2, R version 4.5, on htseq-count output described above. Full software versions for R sessions are listed in RMarkdown output available at github.com/dkbiocode/EmilyEbelRNASeq. The raw counts were normalized and processed according to standard DESeq2 workflows before applying differential expression analysis for all combinations of experimental conditions: temperature 22°C, 26°C or 30°C by mock or WNV infection, resulting in 15 comparisons. Genes were considered differentially expressed in each comparison when the adjusted p-value (Benjamini-Hochberg) was less than 0.1. Interaction formulae were tested but not successful.

### Ortholog mapping of Cx. tarsalis genes

Using the Ctark1 *Cx. tarsalis* assembly [38], coding gene predictions were used in the eggnog/emapper pipeline [78] version 2.1.15 to assign orthologs to the closest annotated species in class Insecta. 11,598 *Cx. tarsalis* genes mapped to *Cx. quinquefasciatus*, while 1,375 mapped to other mosquitos, with 106 mapping to other insects. 1,648 genes *Cx. tarsalis* genes remained unassigned to an ortholog. These mappings were combined with a previous mapping [46], adding 3,007 additional assignments to *Cx. quinquefasciatus*.

This work used the Alpine High-Performance Computing resource at the University of Colorado Boulder. Alpine is jointly funded by the University of Colorado Boulder, the University of Colorado Anschutz, Colorado State University, and the National Science Foundation (award 2201538) [79].

The *Cx. quinquefasciatus* ortholog assignments were also linked to StringDB [80] 12.0 IDs for use in web tools, such as shinyGO. 100% of the StringDB 12.0 IDs were recognized by shinyGO.

### GO term enrichment analysis

GO term enrichment was performed using two methods. The first was executed solely in R with the topGO [81] package 2.60.1 with the classic fisher and pruning algorithms, using GO terms assigned directly from the eggnog/emapper workflow described above. In any given comparison, all genes tested served as the background set, whereas all genes “significant” (adjusted p-value <0.1), were used as the foreground set. Details are available in the repository at 04_DEseq2/GO.Rmd. These procedures yielded no enriched terms after multiple testing correction. To compare with previous analyses, the Culex quinquefasciatus ortholog assignments were submitted to shinyGO 0.86.0, yielding only two related terms for one comparison: Inorganic anion transmembrane transporter activity and Inorganic anion transport for WNV infection at 22°C vs mock infection at 26°C. Gene sets were defined as above and are available in the repository at 04_DEseq2/culex_q_mapping/for_shinyGO.

### AI usage disclosure

Claude Sonnet and GPT Sol 5.6 were used for coding support and pipeline development.

## ACKNOWLEDGEMENTS

This work was supported by funding to Verena (viralemergence.org) from the U.S. National Science Foundation, including NSF BII 2021909 and NSF BII 2213854, and funding from the US National Institutes of Health, National Institute of Allergy and Infectious Diseases (R01-AI067380, R01-AI173206). EAF was supported by training grant 5T32AI162691, and by Colorado State University’s Office of the Vice President for Research’s “Accelerating Innovations in Pandemic Disease” initiative, made possible through support from The Anschutz Foundation. We thank the Cancer Center Genomics Core at the University of Colorado Anschutz Medical Campus and the Cancer Center Support Grant (P30CA046934).

## AUTHOR CONTRIBUTIONS

ENG – investigation, formal analysis, writing – original draft, writing – review & editing

DCK – formal analysis

BM – investigation

EAF – formal analysis

HAO – formal analysis

KXK – investigation

NE – investigation

CLC – resources

GDE – conceptualization, funding acquisition, writing – review & editing

