## Supplemental Materials for "Temperature alters *Culex tarsalis* West Nile virus vector competence, tissue bottlenecks, and transcriptional responses"

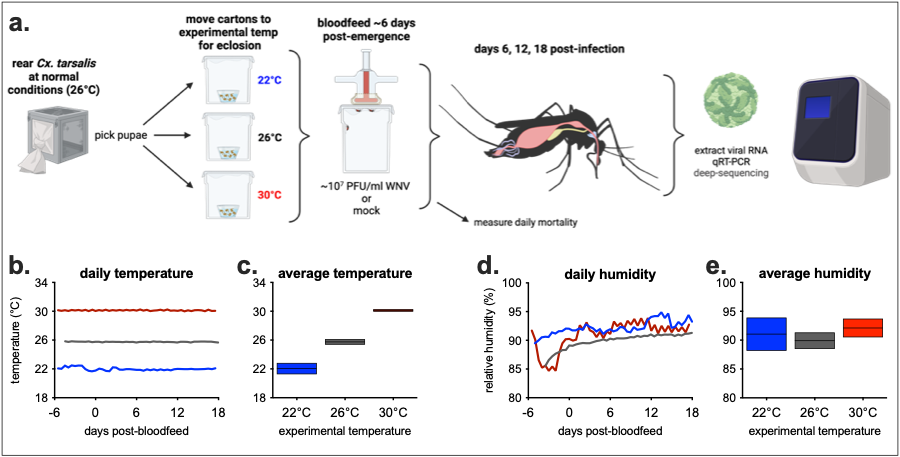

**Supplemental Figure 1. Experimental overview and daily temperature and humidity.** **a)** *Cx. tarsalis* were reared at their normal conditions, then pupae were moved to experimental temperatures. Mosquitoes were given a mock, or WNV-containing bloodmeal, and returned to experimental temperatures. Mortality was measured daily, and on days 6, 12 and 18, a group of mosquitoes was processed for subsequent analyses. **b, d)** Temperature and humidity were measured every hour, and values were averaged over a 12-hour period. **c, e)** Average temperature/humidity ± standard deviation of all hourly measurements over the course of the experiment.

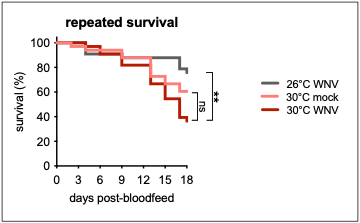

**Supplemental Figure 2. Repeated survival of *Cx. tarsalis* at 30°C.** Survival was measured daily for mock and WNV-infected mosquitoes at 26°C and 30°C (n=33/condition). Log-rank (Mantel-Cox), ** p<0.005.

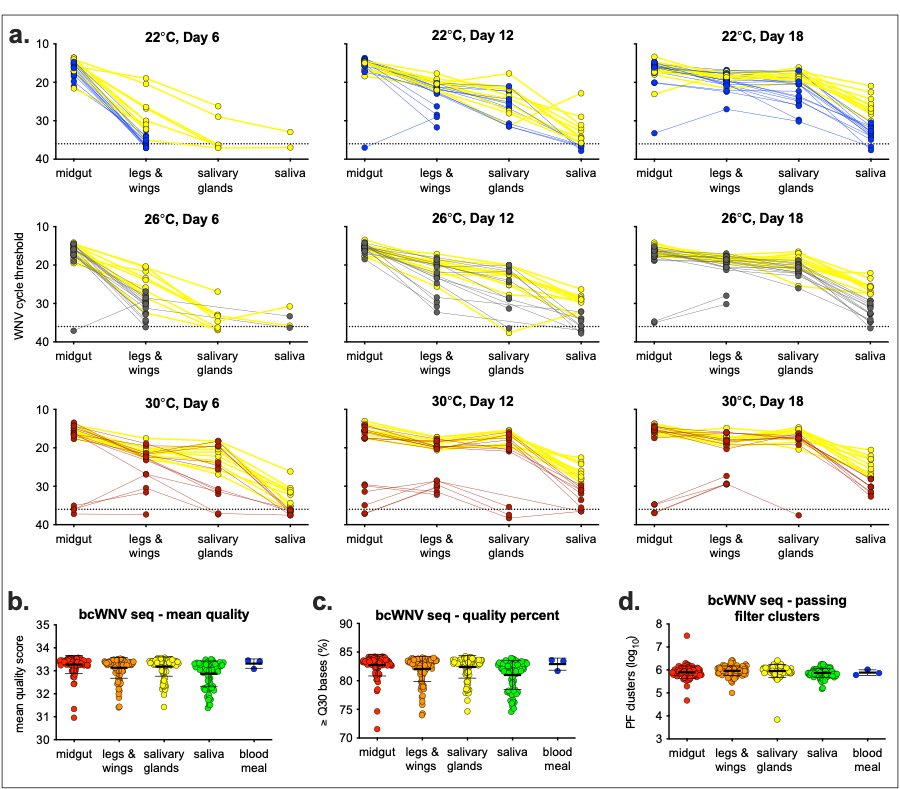

**Supplemental Figure 3. Mosquito barcode deep-sequencing selection and quality control.** **a)** WNV cycle threshold for mosquito samples at each of the three time points (6, 12 and 18 days post-bloodfeed) and temperatures (22°C, 26°C and 30°C). Samples in yellow were selected for barcode deep-sequencing. **b)** Mean quality score, **c)** percent of bases with a quality score ≥ Q30 and **d**) number of clusters passing filter.

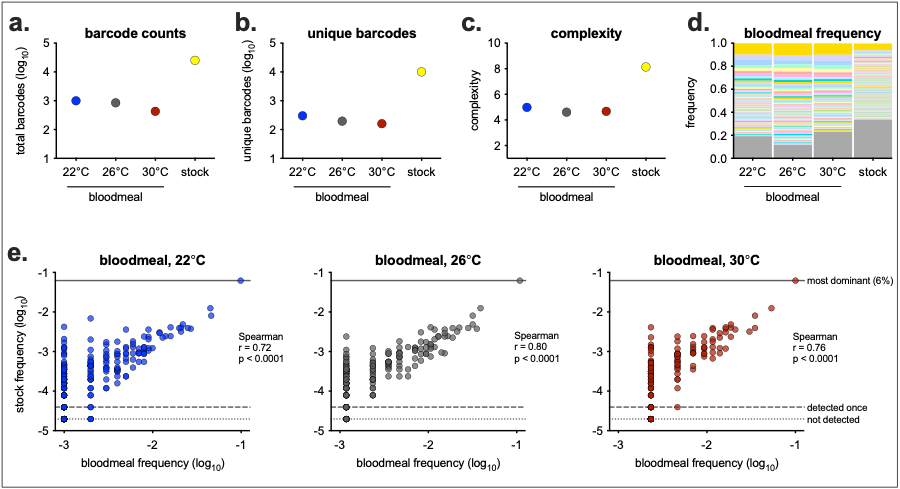

**Supplemental Figure 4. Bloodmeal and virus stock deep-sequencing results.** **a)** Total barcodes, **b)** unique barcodes and **c)** complexity of each of the three experimental temperature bloodmeals (22°C, 26°C and 30°C) and stock virus. **d)** Frequency of each barcode present in the bloodmeals and stocks sorted by most dominant (colors do not correspond to specific barcodes across samples). Grey at the bottom of each bar are barcodes only detected one time in that sample. **e)** The frequency of barcodes in each of the bloodmeals compared to the frequency of that barcode present in the stock. Solid line represents the most dominant barcode in the stock (6% of total reads), dashed line represents the frequency of barcodes only detected one time (0.004%) and dotted line represents barcodes not detected in stock (1/2 the frequency of those detected one time, 0.002%).

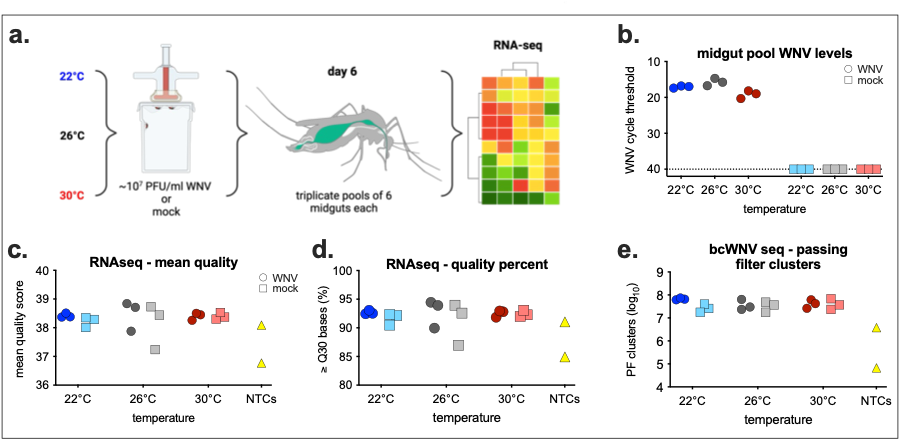

**Supplemental Figure 5. RNAseq experimental overview and sequencing results.** **a)** *Cx. tarsalis* were reared and infected as described previously (**SFig 1a**). At 6 days post-bloodfeed, three pools of 6 midguts each were collected from each experimental condition (WNV and mock, 22°C, 26°C and 30°C). **b)** WNV viral RNA was quantified in each of the pools using qRT-PCR (dashed line is limit of detection). All mosquito pools were processed for host RNA sequencing. **c)** Mean quality score, **d)** percent of bases with a quality score ≥ Q30 and **e**) number of clusters passing filter. NTC = no template controls.

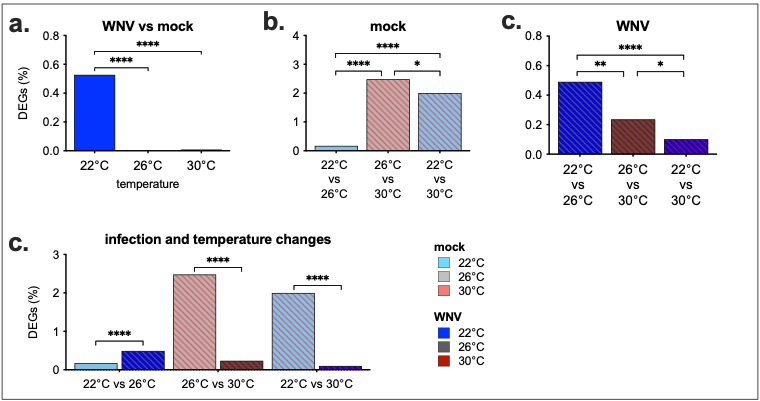
**Supplemental Figure 6. Comparison of differential gene expression by experimental conditions.** **a)** Comparison of DEGs at each experimental temperature in response to WNV infection. Within **b)** mock or **c)** WNV-infected mosquitoes, the impact of changing temperature on DEGs. **d)** Comparing the impact of infection compared to mock, within temperature changes. DEGs are defined as those with an adjusted p-value < 0.1 (shown in **Figure 4a/b/c**). Percent DEGs calculated out of 11,001 total detected genes. Statistics shows Fisher’s exact test comparing DEGs between conditions (* p < 0.05, ** p < 0.005, **** p < 0.0001).

**Supplemental Table 1**. qRT-PCR primer probe sequences

| **Primer** | **Sequence (5’ 🡪 3’)** |
| --- | --- |
| WNV - forward | TCAGCGATCTCTCCACCAAAG |
| WNV - reverse | GGGTCAGCACGTTTGTCATTG |
| WNV - probe | FAM-TGCCCGACCATGGGAGAAGCTC |
| gene1627 - forward | TTGAGGAATTCGGCGTCAT |
| gene1627 - reverse | AGAATCAGTGGGATCGCTTC |
| gene1627 - probe | FAM-AGATGGAGTGGATGCTGCCGAC |
| gene5743 - forward | GGGCGTCAGTTAGCAGTAAT |
| gene5743 - reverse | CGAAGTGATCGGTTTGGTTATG |
| gene5743 - probe | FAM-TCAGCCACCATCAAGGAGAACACG |
| RPL37 – forward | CGTACCTACAACTGGTCGAT |
| RPL37 – reverse | AAACCGTTGCGGAACTTG |
| RPL37 – probe | FAM-CAAGGCCAAGAGGAGGAAGACCAC |

**Supplemental Table 2**. Deep sequencing primers

| **Primer** | **Sequence (5’ 🡪 3’)** |
| --- | --- |
| cDNA | GTCTCGTGGGCTCGGAGATGTGTATAAGAGACAGNNNNNNNNNCAGTGCCATCCACTACAGCGTTCT |
| 1^st^ round PCR  5'-ID-For | TCGTCGGCAGCGTCAGATGTGTATAAGAGACAGNNNNTTCCCCTTCGTCGATGTTGG |
| 1^st^ round PCR  5'-ID-Rev | GTCTCGTGGGCTCGGAGATGTGTAT |
| 2^nd^ round PCR  i5-For | AATGATACGGCGACCACCGAGATCTACAC[i5]TCGTCGGCAGCGTC |
| 2^nd^ round PCR  i7-Rev | CAAGCAGAAGACGGCATACGAGAT[i7]GTCTCGTGGGCTCGG |

**Supplemental Table 3**. Barcode WNV i5 and i7 index primer sequences

| **Primer** | **Sequence (5’ 🡪 3’)** |
| --- | --- |
| i7001 | CAAGCAGAAGACGGCATACGAGATCGCTCAGTTCGTCTCGTGGGCTCGG |
| i7002 | CAAGCAGAAGACGGCATACGAGATTATCTGACCTGTCTCGTGGGCTCGG |
| i7003 | CAAGCAGAAGACGGCATACGAGATATATGAGACGGTCTCGTGGGCTCGG |
| i7004 | CAAGCAGAAGACGGCATACGAGATCTTATGGAATGTCTCGTGGGCTCGG |
| i7005 | CAAGCAGAAGACGGCATACGAGATTAATCTCGTCGTCTCGTGGGCTCGG |
| i7006 | CAAGCAGAAGACGGCATACGAGATGCGCGATGTTGTCTCGTGGGCTCGG |
| i7007 | CAAGCAGAAGACGGCATACGAGATAGAGCACTAGGTCTCGTGGGCTCGG |
| i7008 | CAAGCAGAAGACGGCATACGAGATTGCCTTGATCGTCTCGTGGGCTCGG |
| i7009 | CAAGCAGAAGACGGCATACGAGATCTACTCAGTCGTCTCGTGGGCTCGG |
| i7010 | CAAGCAGAAGACGGCATACGAGATTCGTCTGACTGTCTCGTGGGCTCGG |
| i7011 | CAAGCAGAAGACGGCATACGAGATGAACATACGGGTCTCGTGGGCTCGG |
| i7012 | CAAGCAGAAGACGGCATACGAGATCCTATGACTCGTCTCGTGGGCTCGG |
| i5001 | AATGATACGGCGACCACCGAGATCTACACTCGTGGAGCGTCGTCGGCAGCGTC |
| i5002 | AATGATACGGCGACCACCGAGATCTACACCTACAAGATATCGTCGGCAGCGTC |
| i5003 | AATGATACGGCGACCACCGAGATCTACACTATAGTAGCTTCGTCGGCAGCGTC |
| i5004 | AATGATACGGCGACCACCGAGATCTACACTGCCTGGTGGTCGTCGGCAGCGTC |
| i5005 | AATGATACGGCGACCACCGAGATCTACACACATTATCCTTCGTCGGCAGCGTC |
| i5006 | AATGATACGGCGACCACCGAGATCTACACGTCCACTTGTTCGTCGGCAGCGTC |
| i5007 | AATGATACGGCGACCACCGAGATCTACACTGGAACAGTATCGTCGGCAGCGTC |
| i5008 | AATGATACGGCGACCACCGAGATCTACACCCTTGTTAATTCGTCGGCAGCGTC |
| i5009 | AATGATACGGCGACCACCGAGATCTACACGTTGATAGTGTCGTCGGCAGCGTC |
| i5010 | AATGATACGGCGACCACCGAGATCTACACACCAGCGACATCGTCGGCAGCGTC |
| i5011 | AATGATACGGCGACCACCGAGATCTACACCATACACTGTTCGTCGGCAGCGTC |
| i5012 | AATGATACGGCGACCACCGAGATCTACACGTGTGGCGCTTCGTCGGCAGCGTC |
| i5013 | AATGATACGGCGACCACCGAGATCTACACATCACGAAGGTCGTCGGCAGCGTC |
| i5014 | AATGATACGGCGACCACCGAGATCTACACCGGCTCTACTTCGTCGGCAGCGTC |
| i5015 | AATGATACGGCGACCACCGAGATCTACACGAATGCACGATCGTCGGCAGCGTC |
| i5016 | AATGATACGGCGACCACCGAGATCTACACAAGACTATAGTCGTCGGCAGCGTC |
| i5017 | AATGATACGGCGACCACCGAGATCTACACTCGGCAGCAATCGTCGGCAGCGTC |
| i5018 | AATGATACGGCGACCACCGAGATCTACACCTAATGATGGTCGTCGGCAGCGTC |
| i5019 | AATGATACGGCGACCACCGAGATCTACACGGTTGCCTCTTCGTCGGCAGCGTC |
| i5020 | AATGATACGGCGACCACCGAGATCTACACCGCACATGGCTCGTCGGCAGCGTC |
| i5021 | AATGATACGGCGACCACCGAGATCTACACGGCCTGTCCTTCGTCGGCAGCGTC |
| i5022 | AATGATACGGCGACCACCGAGATCTACACCTGTGTTAGGTCGTCGGCAGCGTC |
| i5023 | AATGATACGGCGACCACCGAGATCTACACTAAGGAACGTTCGTCGGCAGCGTC |
| i5024 | AATGATACGGCGACCACCGAGATCTACACCTAACTGTAATCGTCGGCAGCGTC |
| i5025 | AATGATACGGCGACCACCGAGATCTACACGGCGAGATGGTCGTCGGCAGCGTC |
| i5026 | AATGATACGGCGACCACCGAGATCTACACAATAGAGCAATCGTCGGCAGCGTC |
| i5027 | AATGATACGGCGACCACCGAGATCTACACTCAATCCATTTCGTCGGCAGCGTC |
| i5028 | AATGATACGGCGACCACCGAGATCTACACTCGTATGCGGTCGTCGGCAGCGTC |
| i5029 | AATGATACGGCGACCACCGAGATCTACACTCCGACCTCGTCGTCGGCAGCGTC |
| i5030 | AATGATACGGCGACCACCGAGATCTACACCTTATGGAATTCGTCGGCAGCGTC |
| i5031 | AATGATACGGCGACCACCGAGATCTACACGCTTACGGACTCGTCGGCAGCGTC |
| i5032 | AATGATACGGCGACCACCGAGATCTACACGAACATACGGTCGTCGGCAGCGTC |

**Supplemental Table 4**. Mosquito RNAseq i5 and i7 index primer sequences

| **Primer** | **Sequence (5’ 🡪 3’)** |
| --- | --- |
| NEBNext i701 | CAAGCAGAAGACGGCATACGAGATCGAGTAATGTGACTGGAGTTCAGACGTGTGCTCTTCCGATC*T |
| NEBNext i702 | CAAGCAGAAGACGGCATACGAGATTCTCCGGAGTGACTGGAGTTCAGACGTGTGCTCTTCCGATC*T |
| NEBNext i703 | CAAGCAGAAGACGGCATACGAGATAATGAGCGGTGACTGGAGTTCAGACGTGTGCTCTTCCGATC*T |
| NEBNext i704 | CAAGCAGAAGACGGCATACGAGATGGAATCTCGTGACTGGAGTTCAGACGTGTGCTCTTCCGATC*T |
| NEBNext i705 | CAAGCAGAAGACGGCATACGAGATTTCTGAATGTGACTGGAGTTCAGACGTGTGCTCTTCCGATC*T |
| NEBNext i501 | AATGATACGGCGACCACCGAGATCTACACTATAGCCTACACTCTTTCCCTACACGACGCTCTTCCGATC*T |
| NEBNext i502 | AATGATACGGCGACCACCGAGATCTACACATAGAGGCACACTCTTTCCCTACACGACGCTCTTCCGATC*T |
| NEBNext i503 | AATGATACGGCGACCACCGAGATCTACACCCTATCCTACACTCTTTCCCTACACGACGCTCTTCCGATC*T |
| NEBNext i504 | AATGATACGGCGACCACCGAGATCTACACGGCTCTGAACACTCTTTCCCTACACGACGCTCTTCCGATC*T |

**Supplemental Table 5**. *Cx. tarsalis* gene ID and name

| **gene ID** | **gene name** |
| --- | --- |
| gene1501 | Dome |
| gene6092 | Cactus |
| gene10322 | MyD88 |
| gene11340 | Imd |
| gene5564 | DefensinA |
| gene3946 | NFKB1 |
| gene3044 | Ago2 |
| gene6258 | R2D2 |
| gene8050 | Hsp70 |
| gene54 | HSP90AB1 |
| gene55 | HSP90AB1* |
| gene6002 | RPL37 |
